# *Cucumeropsis mannii* seed oil (CMSO) ameliorates adipokines dysfunction and dyslipidemia in male Wistar rats exposed to Bisphenol-A

**DOI:** 10.1101/2022.01.04.474972

**Authors:** P. M. Aja, D. A Chukwu, P.C. Agu, O. G. Ani, E. U. Ekpono, H. A. Ogwoni, J. N. Awoke, Patience N. Ogbu, L. Aja, F. E. Nwite, O. U. Ukachi, O. U. Orji, P. C. Nweke, C. O. Egwu, E. U. Ekpono, G. O. Ewa, I. O. Igwenyi, E. U. Alum, D. E. Uti, Deusdedit Tusubira, C. E. Offor, A. Njoku, E. K Maduagwuna

## Abstract

Bisphenol-A (BPA) and its analog are extensively utilized in the production of plastics which are rather ubiquitous in our environment. At high temperatures, BPA is leached into water and food packed in plastic containers. This research investigated the ameliorative effects of CMSO on adipokines dysfunction and dyslipidemia in male Wistar rats exposed to Bisphenol-A. thirty-six (36) albino rats weighing 100 - 200 g were randomly assigned into six (6) different experimental groups of controls (1, 2, and 3) and the tests (4, 5, and 6). Group 1 was given only 1 ml of olive oil, group 2 received 100 mg/Kg body weight (b.w) of BPA, group 3 was given 7.5 ml/Kg b.w of CMSO, groups 4, 5, and 6 received 100 mg/Kg b.w of BPA and 7.5, 5 and 2.5 mg/Kg b.w of CMSO respectively. CMSO and BPA were concurrently administered via oral intubation for periods of 42 days. Lipid profile and adipokines levels were determined in plasma and adipose tissue. BPA in male rats significantly (p<0.05) elevated the levels of cholesterol, triglycerides, LDL-C, liptin, and coronary and atherogenic risk indices in plasma and adipose tissue with reductions in HDL-C and adiponectin levels. BPA plus CMSO in male rats significantly (p<0.05) decreased the levels of cholesterol, triglycerides, LDL-C, liptin, and coronary and atherogenic risk indices with an elevation of HDL-C and adiponectin levels in both plasma and adipose tissue. These results suggest that CMSO could be useful in the management of cardiovascular-related diseases induced by BPA.

## 1. Introduction

An array of agents dubbed “endocrine-disrupting chemicals” (EDCs) that function by mimicking the endogenous hormones hamper the signaling system, thereby orchestrating numerous metabolic disorders leading to an array of ailments (Flint *et al*., 2012; Preethi *et al*., 2014; Metz, 2016). One of such prominent xenobiotics that functions as an endocrine disruptor is 2,2-Bis(4-hydroxy-phenyl) propane (Šutiaková, 2012; Gabr *et al*., 2017), shortened as Bisphenol A (BPA).

BPA is a carbon-based synthetic chemical that is commonly used as a monomer in the production of polycarbonate plastics (electronic devices, kitchen utensils, baby and water bottles, sports equipment, and medical devices) and epoxy resin (Kelley *et al*., 2015; Almeida *et al*., 2018; Talpade *et al*., 2018), like dental sealants (vom Saal and Hughes, 2005), as an antioxidant in polyvinylchloride (PVC) plastics and as an inhibitor of end polymerization in PVC and a variety of other applications (Caporossi and Papaleo, 2015). Ingestion, inhalation, or dermal contact have been identified as the major routes of human exposure to BPA (Talpade *et al*., 2018). It has been widely reported that increased temperature, elongated storage time, repeated usage, and acidic or basic food or beverages in cans and polycarbonate plastics can facilitate the leaching of the BPA into the contents (vom Saal and Hughes, 2005; Šutiaková, 2012; Adeghe and Emejulu, 2016). The leached BPA ultimately hydrolyze in the food and liquids making the dietary consumption the main source of human exposure (Šutiaková, 2012; Adeghe and Emejulu, 2016). Similarly, environmental pollution *inter alia* may lead to exacerbated exposure of BPA (Stragierowicz, 2015). One of the crucial endocrine organs in the biological system is the adipose tissue that produces peptide hormones, known as adipokines (liptin and adiponectin) that may act locally (autocrine or paracrine) or systemically (endocrine action), carrying information about the adequacy of the energy reserves (Triacylglycerides, TAG) stores in the adipose tissue to other tissues and the brain (Nelson and Cox, 2008). In addition to this, the adipose tissue also plays a central function as fuel depot as it houses TAGs, as well as maintaining the temperature homeostasis in the animals (Müllerová and Kopecký, 2007). Like numerous other EDCs, BPA is lipophilic with long half-life, hence it has been reported to bioaccumulate in the adipose tissue (Doke *et al*., 2018; Akash *et al*., 2020). Consequently, predisposing the subjects to weight gain as the BPA has been reported to induce adipogenesis (Ohlstein *et al*., 2014; Ariemma *et al*., 2016; Yang *et al*., 2016) even in children (Menale *et al*., 2015; Desai *et al*., 2018). Ultimately, the BPA effect on the adipose suggests that it’s an obesogen – elicits abnormal fats accumulation leading to obesity (vom Saal *et al*., 2012; Ariemma *et al*., 2016; Apau *et al*., 2018). The functioning of the adipose tissue’s adipokines has also been reported to be marred by the BPA exposure (Ben-jonathan *et al*., 2009; Rönn *et al*., 2014). In the light of this, therefore, BPA exposure being ubiquitous must be accosted by a countermeasure preferably via a dietary source as this also has proven to be the main route of exposure.

Plants in addition to the provision of food to man also afford an array of structurally distinct secondary metabolites known as natural products that have diverse therapeutic potentials contributing to the improvement of health care of humanity (Watal *et al*., 2014). Cucurbitaceae family, sometimes referred to also as the gourd family is reputed to be one of the genetically endowed restorative plants in the plant kingdom (Dhakad *et al*., 2017) coupled with their provision of lipids and proteins in diets Desai *et al*., 2018.

*Cucumeropsis mannii* of the Cucurbitaceae family is a non-hardy legume growing as a tendril climber or creeper in a wet humid climate, particularly in South-western Nigeria, Central, Littoral, and Southern region of Cameroon (Essien *et al*., 2012; Watal *et al*., 2014). Common names for this plant include ‘Egusi-it in Igbo, ‘Egusi’ in Yoruba, and ‘Agushi’ in Hausa (Adeghe *et.al* 2016). In English, it is known as Mann’s cucumeropsis and white-seed melon (Adeghe *et al*., 2016)

Studies have indicated that the *C. mannii* seed oil could be potential good edible oils for reducing cardiovascular illnesses (Amin *et.al*., 2018). It has also been reported that the *C. mannii* seed oil possesses inhibitory abilities on phosphodiesterase– 5 and arginase and thus suggesting their erectogenic potentials (Almeida *et al*., 2018). Therefore, this study was carried out to ascertain whether the *C. mannii* seed oil could have the potentials of maintaining systemic lipid homeostasis and ameliorating the toxic effect of Bisphenol A in the adipose tissues of rats.

## 2. Materials and Methods

### 2.1 Chemicals and Reagents

The Bisphenol A ([2,2-bis(4-hydroxyphenyl)propane] in the form of pure pellets and the reagents used are of analytical grades, majorly obtained from Sigma Aldrich Company, UK through Bristol Scientific.

### 2.2 Collection and Authentication of Plant Material

The plant material used in this study was *Cucumeropsis mannii* Naud Seed which was purchased from market women in Iboko market, Izzi L.G.A., Ebonyi State. The plant was classified and authenticated by a plant taxonomist Mr. Nwankwo of the Department of Applied Biology of Ebonyi State University, Abakaliki, Nigeria. Part of the identified plant was kept in the herbarium of the Applied Biology Department, Ebonyi State University, Abakaliki, Ebonyi State, Nigeria, for reference purposes.

### 2.3 Extraction of *Cucumeropsis mannii* Naud Seed Oil (CMSO)

The *Cucumeropsis mannii* Naud seed oil was extracted using the local method as described by Kate *et al*., (2014).

### 2.4 Acute Toxicity Test

The acute toxicity of *Cucumeropsis mannii* seed oil in male Wistar Albino rats was determined by fixed-dose methods (OECD, 2009).

### 2.5 Animals Handling

The experimental animals used in this study were albino Wister rats purchased from the Animal House of Faculty of Veterinary Medicine, University of Nigeria, Nsukka, Enugu, Nigeria. The rats were kept in stainless steel rats’ cages in a well-ventilated animal house of the Biochemistry Department, Ebonyi State University Abakaliki. They were acclimatized for seven days under good laboratory conditions (12 hours light/dark cycle; room temperature). They were allowed free access to standard rodent chow (Vital feed^®^, Grand Cereals Ltd, Jos, Nigeria) and water *ad libitum*. The procedure for experimental studies was performed consistent with the National Institute of Health Guide for the Care and Use of Laboratory Animals (NIH, 1996). The Department of Biochemistry Ethical Review Committee, Ebonyi State University, Abakaliki, Nigeria, approved this study with ethical no: **EBSU/BCH/ET/20/003.**

### 2.6 Experimental Design

A total of thirty-six (36) male albino Wistar rats were used for this study. The rats were randomly assigned into six (6) experimental groups of 1, II, III, IV, V, and VI with six (6) rats in each group. Groups I, II, and III were control groups while groups IV, V, and VI were the treatment groups.

Group I: Rats received 1 ml of olive only and served as the normal control known to be control group 1 (CG1).
Group II: Rats received 100 mg/Kg body weight(b.w) of BPA orally and served as the BPA control group known to be control group 2 (CG2) (Samova *et al*., 2018).
Group III: Rats received 7.5 ml/Kg b.w of *Cucumeropsis mannii* Naud Seed Oil (CMSO) orally and served as the CMSO control group known to be control group 3 (CG3).
Group IV: Rats were pre-administered 100 mg/Kg b.w of BPA and were treated with 7.5 ml/Kg b.w of CMSO and served as treatment group 1 (TG1).
Group V: Rats were pre-administered 100 mg/Kg b.w of BPA and were treated with 5 ml/kg b.w of CMSO and served as treatment group 2 (TG2).
Group VI: Rats were pre-administered 100 mg/Kg b.w of BPA and were treated with 2.5 ml/Kg b.w of CMSO and served as a treatment group (TG3).

Administration of both BPA and CMSO were concurrently by oral intubation once every day for six weeks.

### 2.7 Tissue Sample Collection

Overnight after the trial, the animals were sacrificed by cervical dislocation under mild anesthesia. The venous blood was collected into plain sample bottles. The blood samples were centrifuged at 3,000 rpm for 5 min and the sera were carefully pipetted into properly labeled tubes for analysis. The adipose tissues of the rats were excised, cleansed of superficial connective tissues, and stored in ice-cold 0.25 M sucrose solution (1:5 w/v).

### 2.8 Determination of Biochemical Parameters

a. **Determination of Lipid Profile of the rats:** The concentration of Triglyceride was determined calorimetrically using the method of Tietz (1990). The method of Bachorik (1996) was used to determine the HDL-cholesterol concentration. The serum concentration of the free fatty acids was determined using the method by Anstall (1965). The plasma total cholesterol concentration was determined using the enzymatic saponification procedure by Allain *et al*. (1974). LDL-C was determined from HDL-C, Total CHOL and TAG using a Friedewald equation (Friedewald *et al*., 1972).
b. **Determination of Atherogenic index (AI) and coronary risk index (CRI) of the rats:** Atherogenic index (AI) and coronary risk index (CRI) of the rats were expressed as ratios of LDL cholesterol to HDL cholesterol and total cholesterol to HDL cholesterol, respectively.
c. **Determination of Liptin Level:** Liptin level was determined according to the methods of Hotta (2000).
d. **Determination of Adiponectin Level:** Adiponectin level was determined according to the method of Yamauchi *et al*. (2001).

### 2.9 Statistical Analysis

Statistical analyses were carried out using a Graph pad prism. Results were expressed as mean ± standard deviation (SD). The various data were subjected to one-way analysis of variance (ANOVA), the difference between the samples was determined by Duncan’s multiple range test setting P values at 0.05.

## 3. Results

### 3.1 Effect of CMSO on serum lipid profile in BPA interference of lipid metabolism in male rats

Administration of BPA significantly (p<0.05) elevated the serum levels of cholesterol, triglycerides, and LDL-C with a reduction in HDL-C level in rats (Figures 1–4). However, co-administration of BPA and CMSO significantly (p<0.05) reduced cholesterol, triglycerides, and LDL-C with an elevation of HDL-C as shown in Figures 1–4.

**Figure 1:**
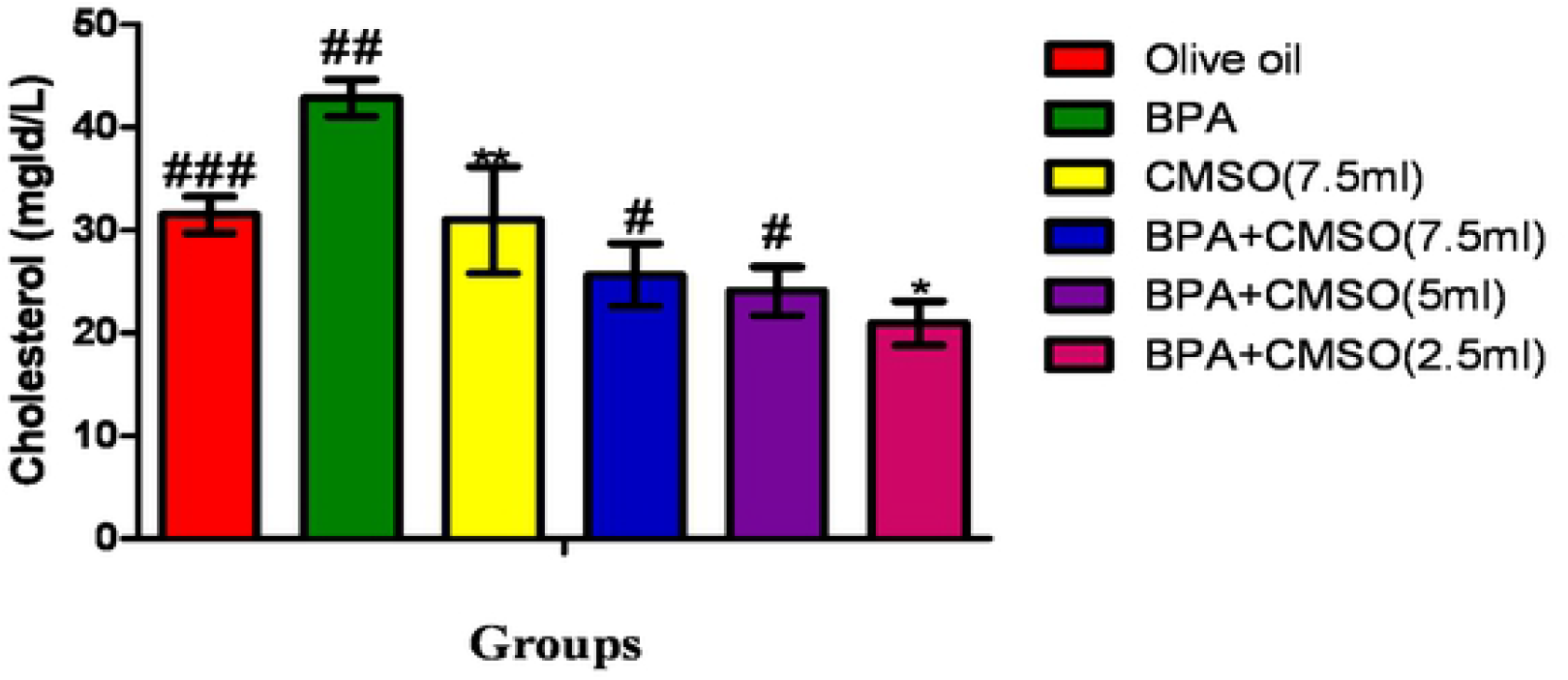
Effect of CMSO on the serum Cholesterol level in BPA interference in lipid metabolism in albino rats. Data are shown as mean ± S.D (n=6). Mean values with the different signs are significantly different at P<0.05. BPA (Bisphenol A), CMSO (*Cucumeropsis mannii* Seed Oil).

**Figure 2:**
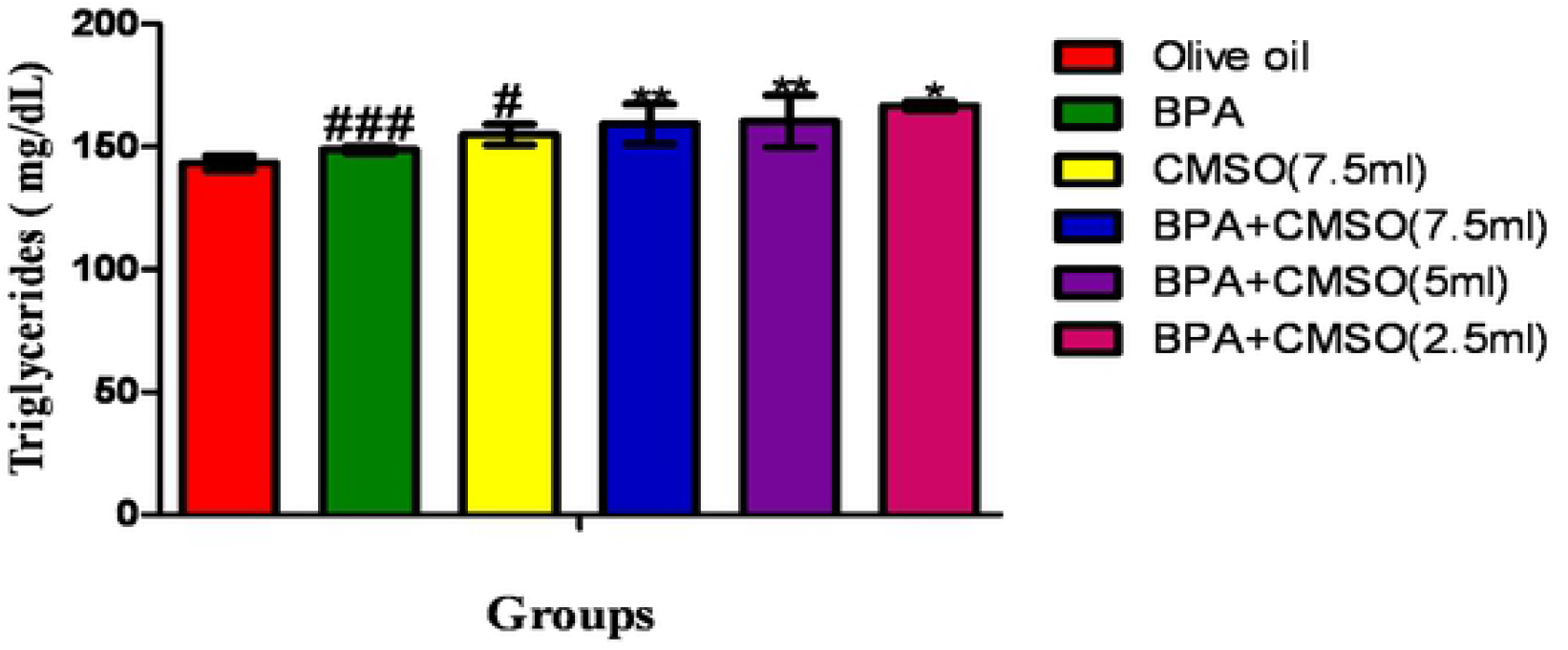
Effect of CMSO on serum triglyceride level in BPA interference in lipid metabolism in albino rats. Data are shown as mean ± S.D (n=6). Mean values with the different signs are significantly different at P<0.05. BPA (Bisphenol A), CMSO (*Cucumeropsis mannii* Seed Oil).

**Figure 3:**
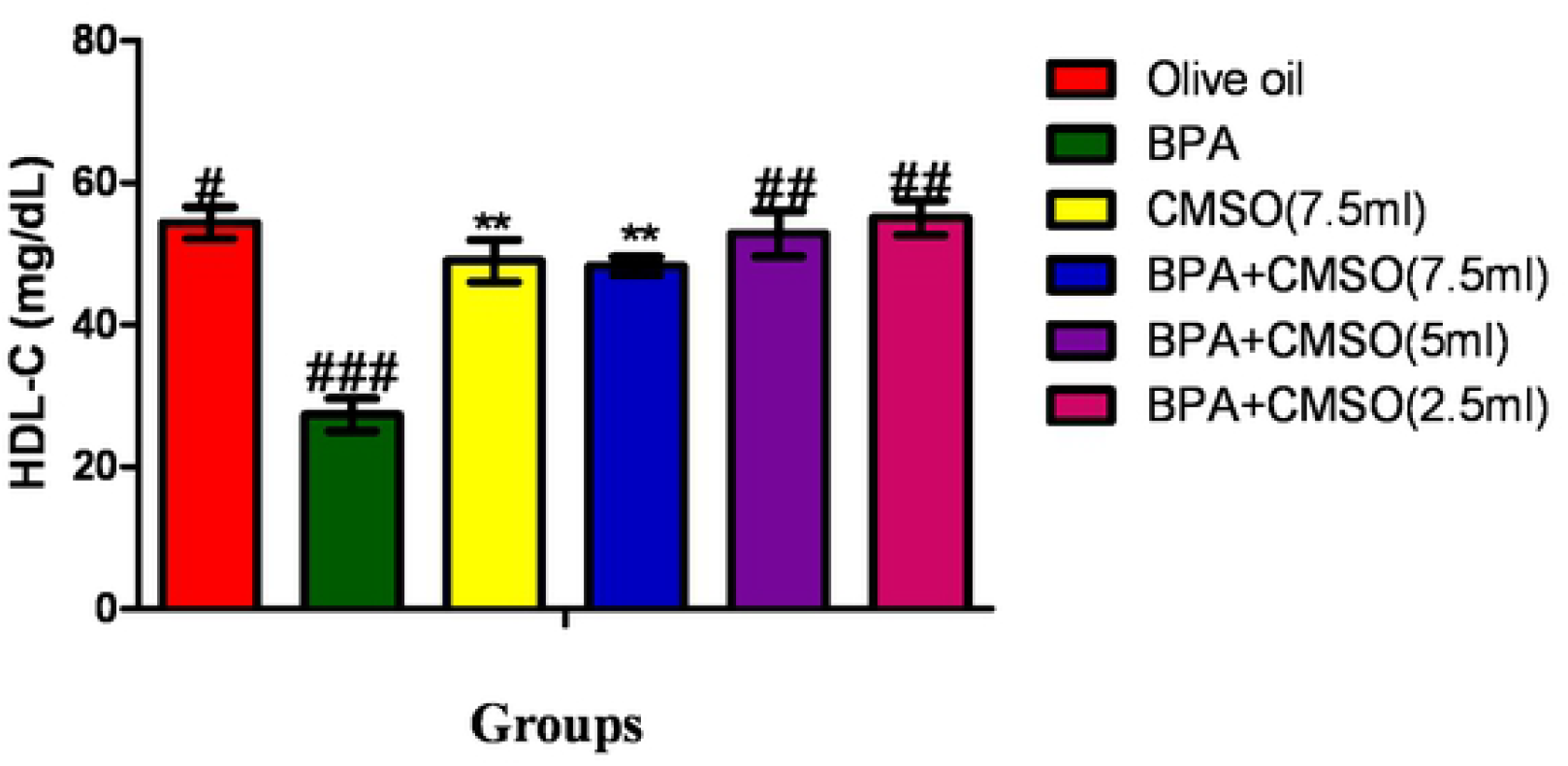
Effect of CMSO on serum HDL-C level in BPA interference in lipid metabolism in albino rats. Data are shown as mean ± S.D (n=6). Mean values with the different signs are significantly different at P<0.05. BPA (Bisphenol A), CMSO (*Cucumeropsis mannii* Seed Oil).

**Figure 4:**
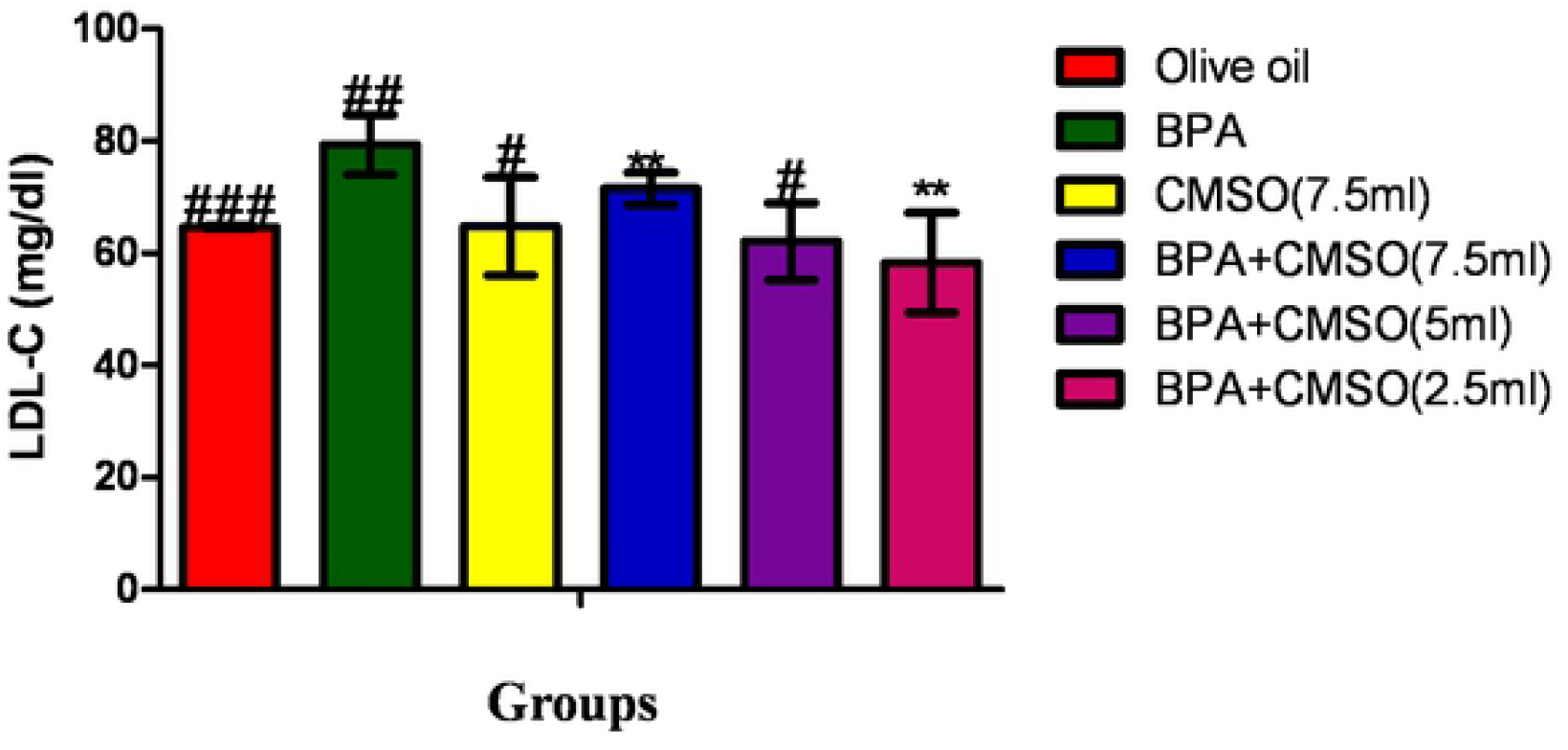
Effect of CMSO on serum LDL-C level in BPA interference in lipid metabolism in albino rats. Data are shown as mean ± S.D (n=6). Mean values with the different signs are significantly different at P<0.05. BPA (Bisphenol A), CMSO (*Cucumeropsis mannii* Seed Oil).

### 3.2 Effect of CMSO on adipose tissue lipid profile in BPA interference of lipid metabolism in male rats

Administration of BPA significantly (p<0.05) elevated the levels of cholesterol, triglycerides, and LDL-C with a reduction in HDL-C level in rat adipose tissue (Figures 1–4). However, co-administration of BPA and CMSO significantly (p<0.05) reduced cholesterol, triglycerides, and LDL-C with the elevation of HDL-C as shown in Figures 5–8.

**Figure 5:**
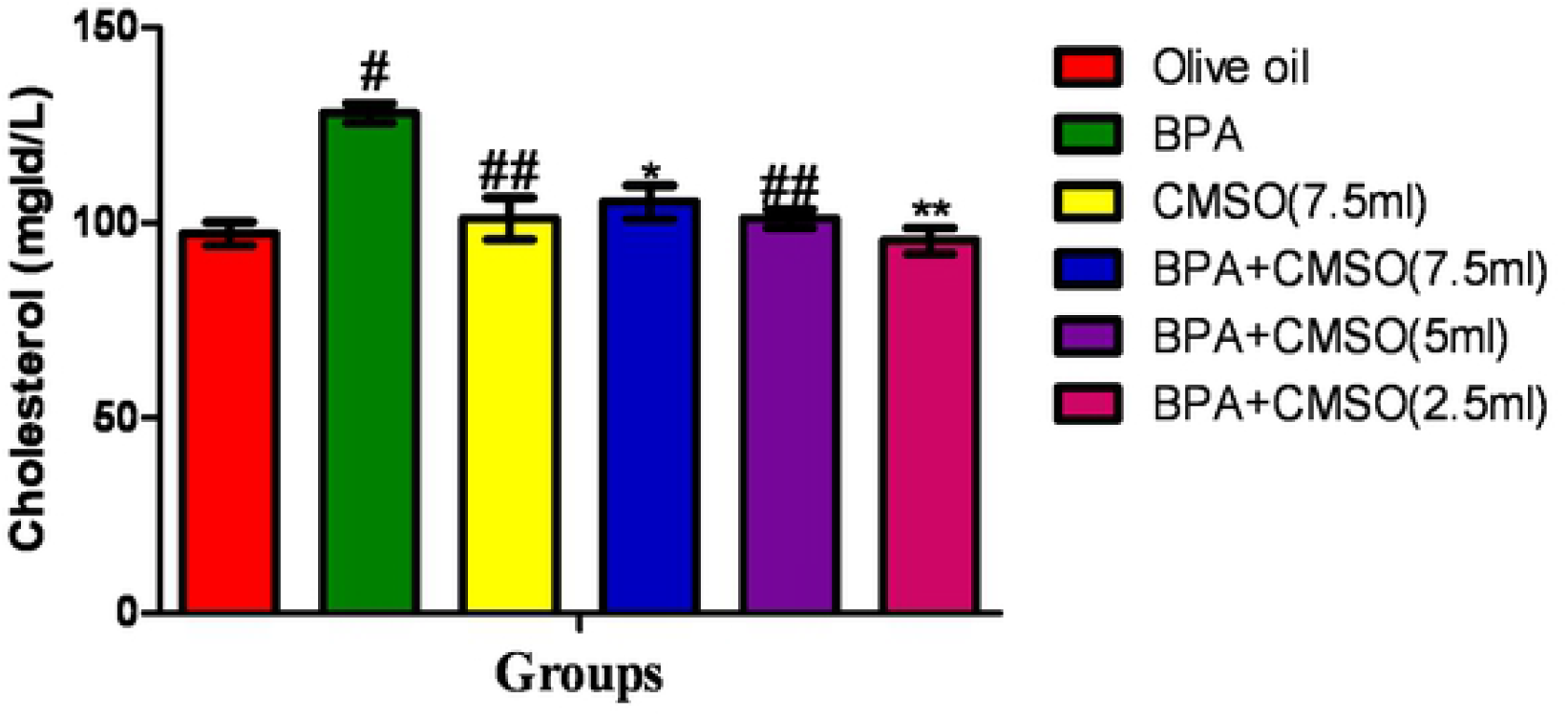
Effect of CMSO on Cholesterol level in adipose tissue in BPA interference in lipid metabolism in albino rats. Data are shown as mean ± S.D (n=6). Mean values with the different signs are significantly different at P<0.05. BPA (Bisphenol A), CMSO (*Cucumeropsis mannii* Seed Oil).

**Figure 6:**
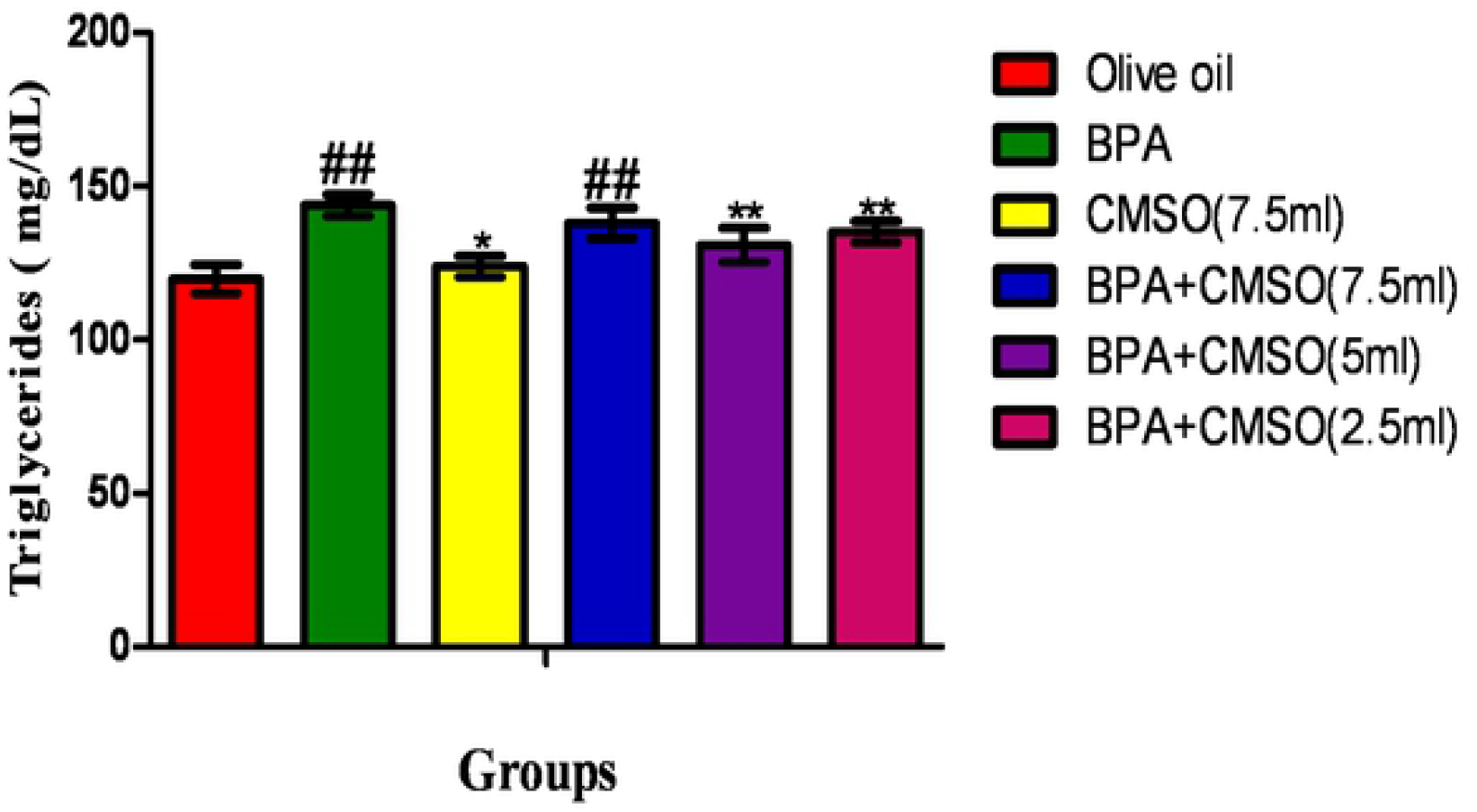
Effect of CMSO on Triglycerides level in adipose tissue in BPA interference in lipid metabolism in albino rats. Data are shown as mean ± S.D (n=6). Mean values with the different signs are significantly different at P<0.05. BPA (Bisphenol A), CMSO (*Cucumeropsis mannii* Seed Oil).

**Figure 7:**
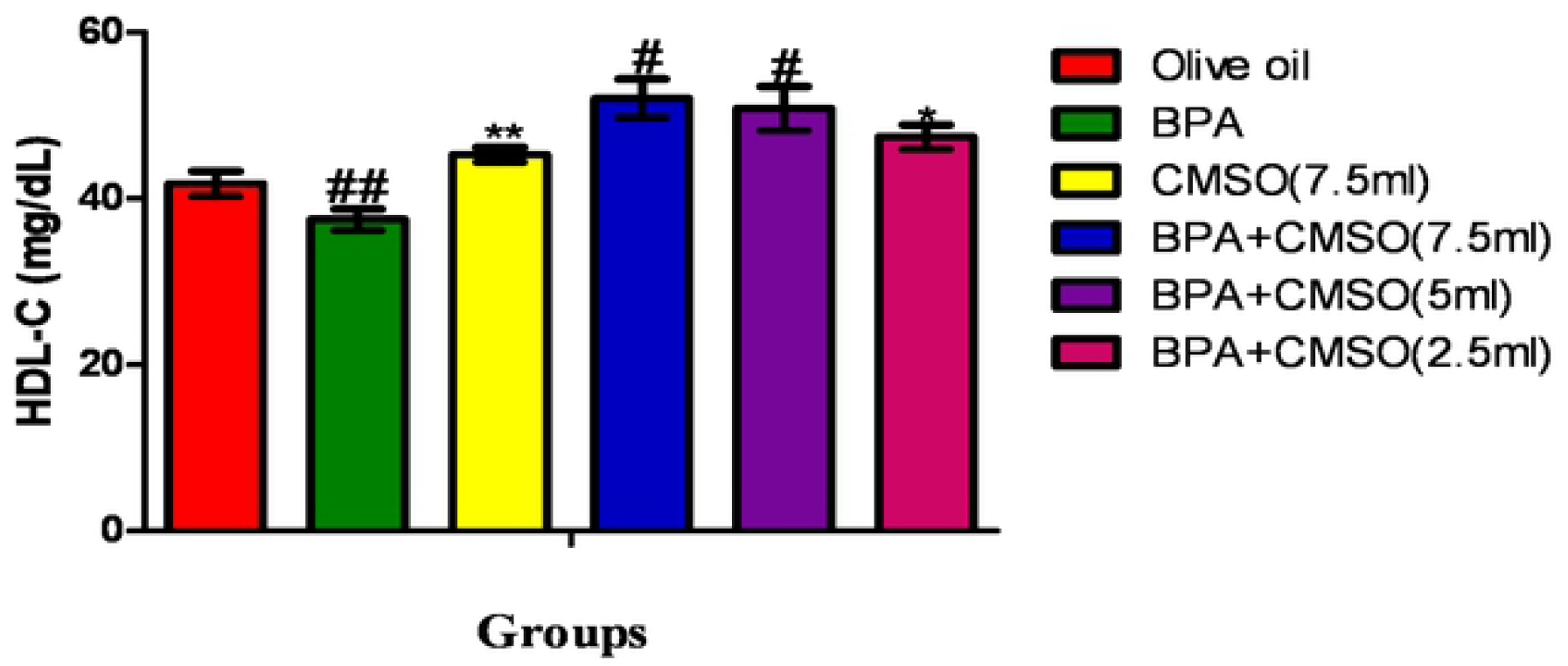
Effect of CMSO on HDL-C level in adipose tissue in BPA interference in lipid metabolism in albino rats. Data are shown as mean ± S.D (n=6). Mean values with the different signs are significantly different at P<0.05. BPA (Bisphenol A), CMSO (*Cucumeropsis mannii* Seed Oil).

**Figure 8:**
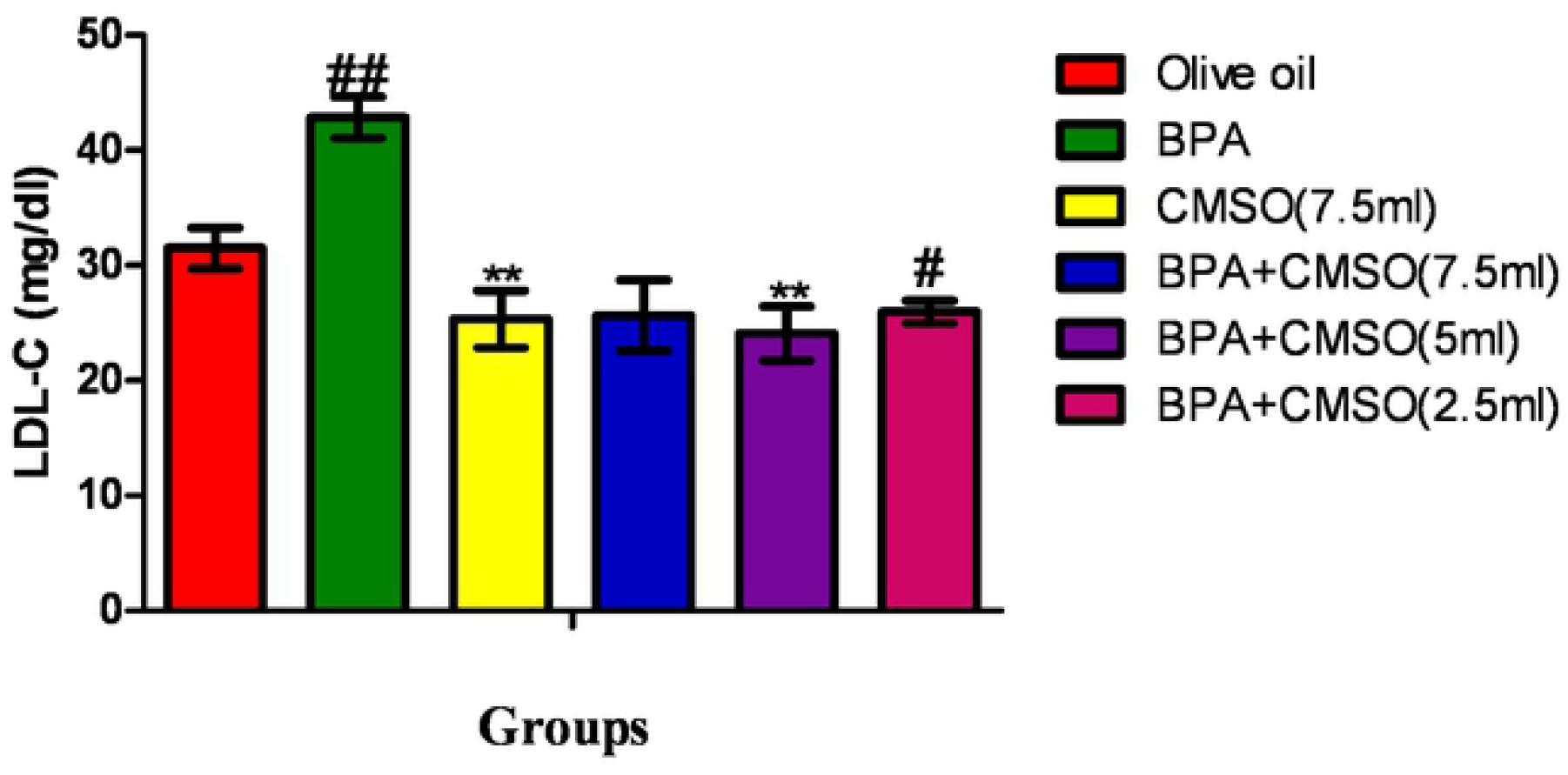
Effect of CMSO on LDL-c level in adipose tissue in BPA interference in lipid metabolism in albino rats. Data are shown as mean ± S.D (n=6). Mean values with the different signs are significantly different at P<0.05. BPA (Bisphenol A), CMSO (*Cucumeropsis mannii* Seed Oil).

### 3.3 Effect of CMSO on Serum Coronary Risk Index in BPA interference of lipid metabolism in male rats

Bisphenol-A (BPA) administration significantly (p<0.05) elevated the coronary risk index in serum and adipose tissue of male rats (Figures 9 and 10). Co-administration of BPA and CMSO in male rats significantly(p<0.05) reduced the coronary risk index in both serum and adipose tissue of rats as shown in Figures 9 and 10.

**Figure 9:**
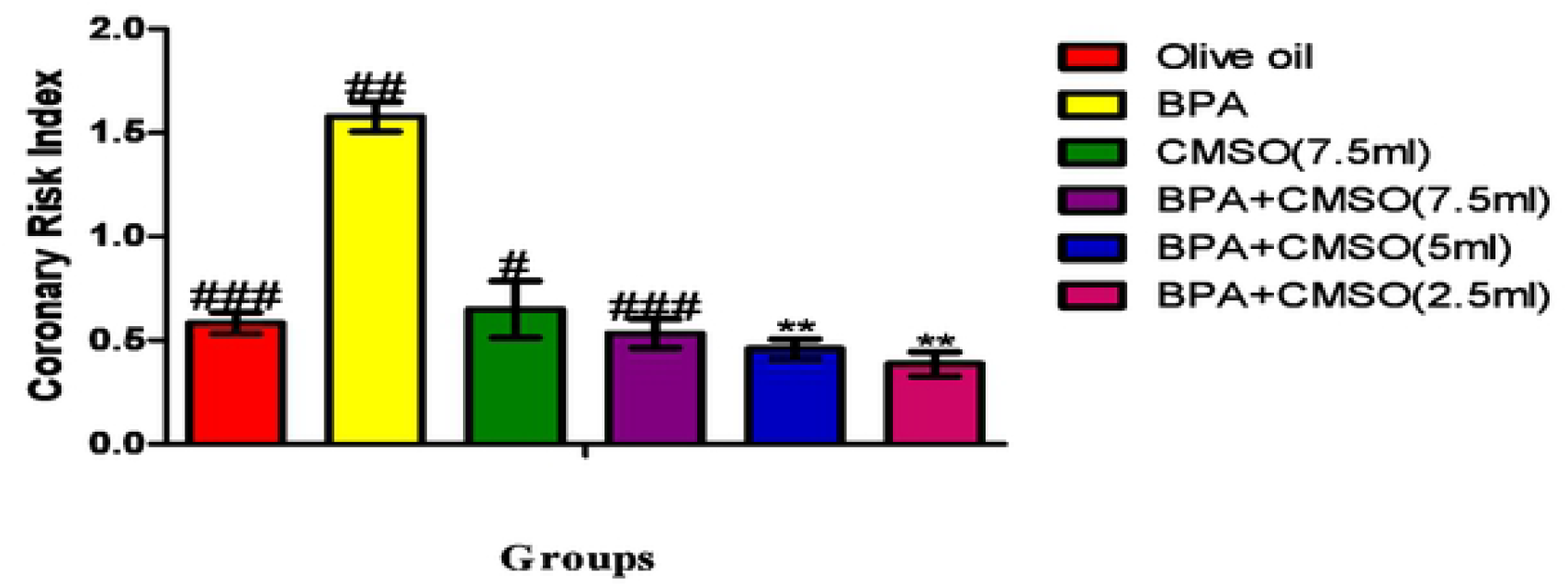
Effect of CMSO on Serum Coronary risk index in BPA interference in lipid metabolism in albino rats. Data are shown as mean ± S.D (n=6). Mean values with the different signs are significantly different at P<0.05. BPA (Bisphenol A), CMSO (*Cucumeropsis mannii* Seed Oil).

**Figure 10:**
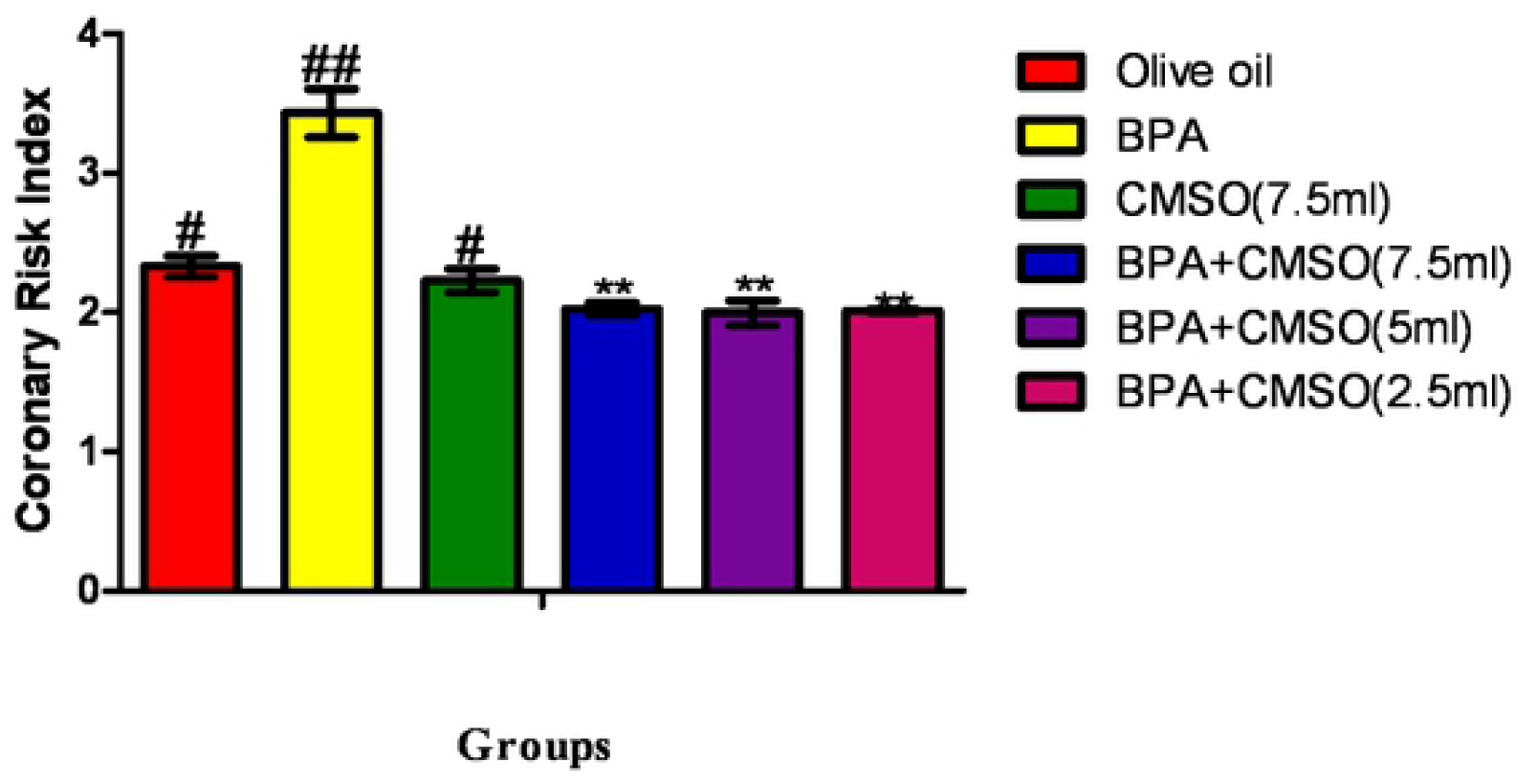
Effect of CMSO on Adipose tissue Coronary risk index in BPA interference in lipid metabolism in albino rats. Data are shown as mean ± S.D (n=6). Mean values with the different signs are significantly different at P<0.05. BPA (Bisphenol A), CMSO (*Cucumeropsis mannii* Seed Oil).

### 3.4 Effect of CMSO on Adipose Tissue Antherogenic Risk Index in BPA interference of lipid metabolism in male rats

Bisphenol-A (BPA) administration significantly (p<0.05) elevated the atherogenic risk index in serum and adipose tissue of male rats (Figures 11 and 12). Co-administration of BPA and CMSO in male rats significantly(p<0.05) reduced the atherogenic risk index in both serum and adipose tissue of rats as shown in Figures 11 and 12.

**Figure 11:**
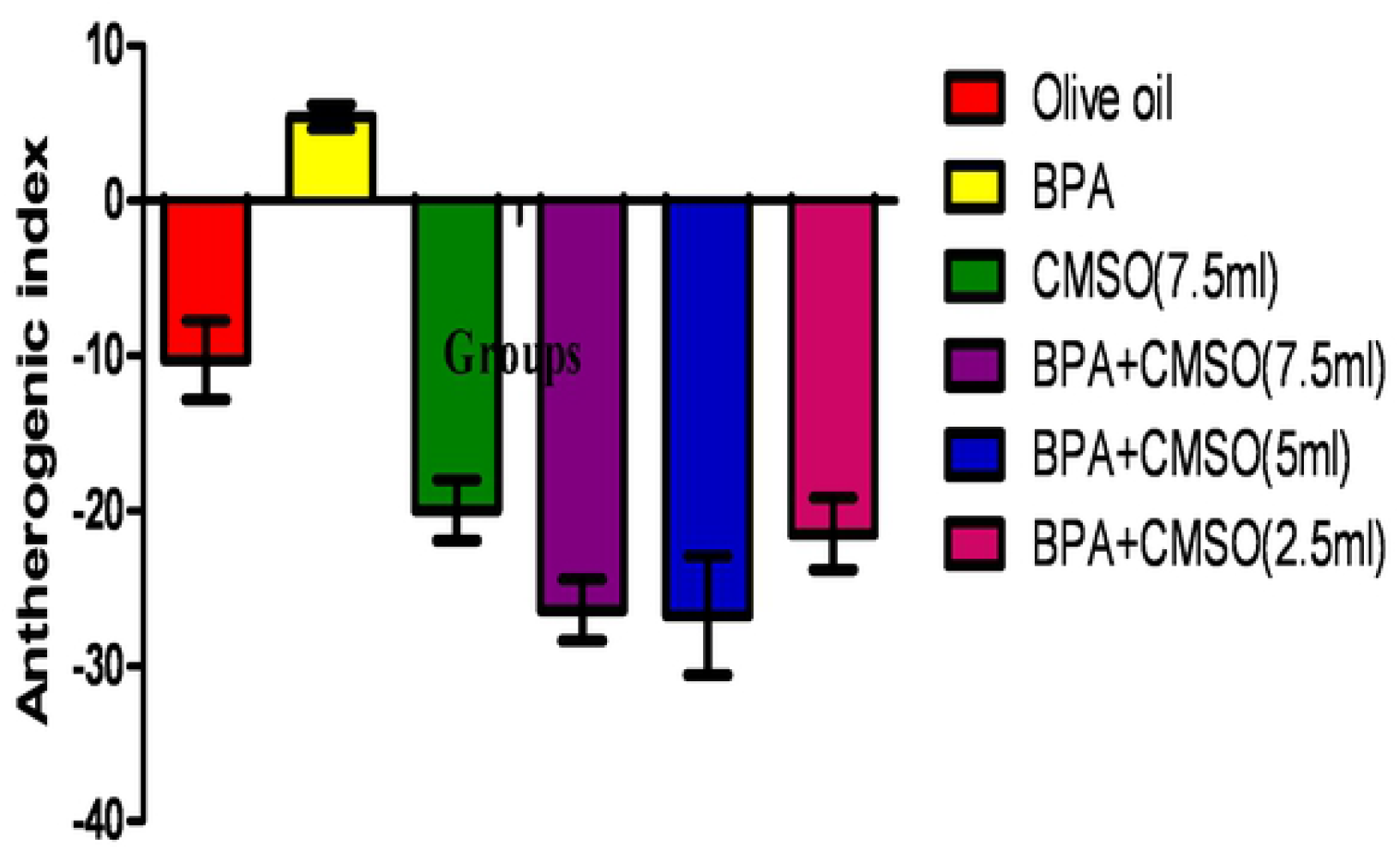
Effect of CMSO on Serum Antherogenic risk index in BPA interference in lipid metabolism in albino rats. Data are shown as mean ± S.D (n=6). Mean values with the different signs are significantly different at P<0.05. BPA (Bisphenol A), CMSO (*Cucumeropsis mannii* Seed Oil).

**Figure 12:**
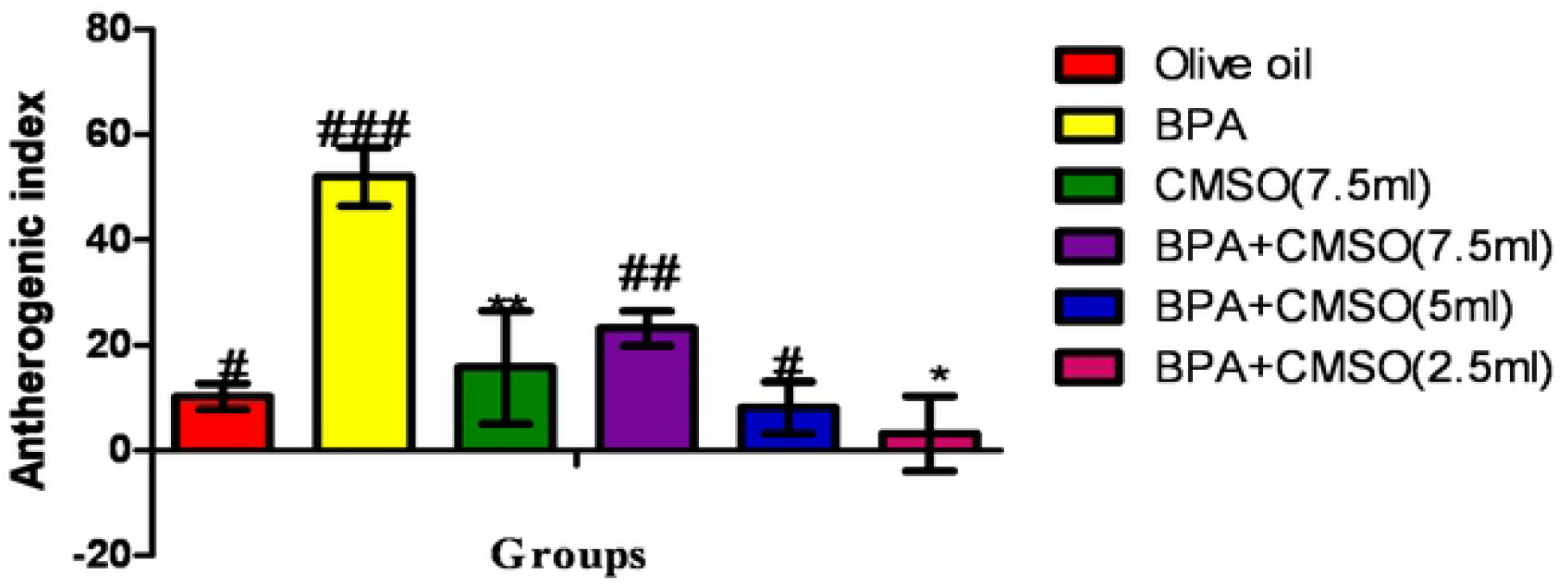
Effect of CMSO on Adipose tissue Antherogenic risk index in BPA interference in lipid metabolism in albino rats. Data are shown as mean ± S.D (n=6). Mean values with the different signs are significantly different at P<0.05. BPA (Bisphenol A), CMSO (*Cucumeropsis mannii* Seed Oil).

### 3.5 Effect of CMSO on Adipose Tissue Liptin and Adiponectin levels in BPA interference of lipid metabolism in male rats

Bisphenol-A (BPA) administration significantly (p<0.05) elevated liptin level and reduced adiponectin level in male rats adipose tissues (Figures 13 and 14. Co-administration of BPA and CMSO in male rats significantly(p<0.05) reduced the level of liptin in a dose-dependent manner and elevated the adiponectin level in male rats as shown in Figures 13 and 14.

**Figure 13:**
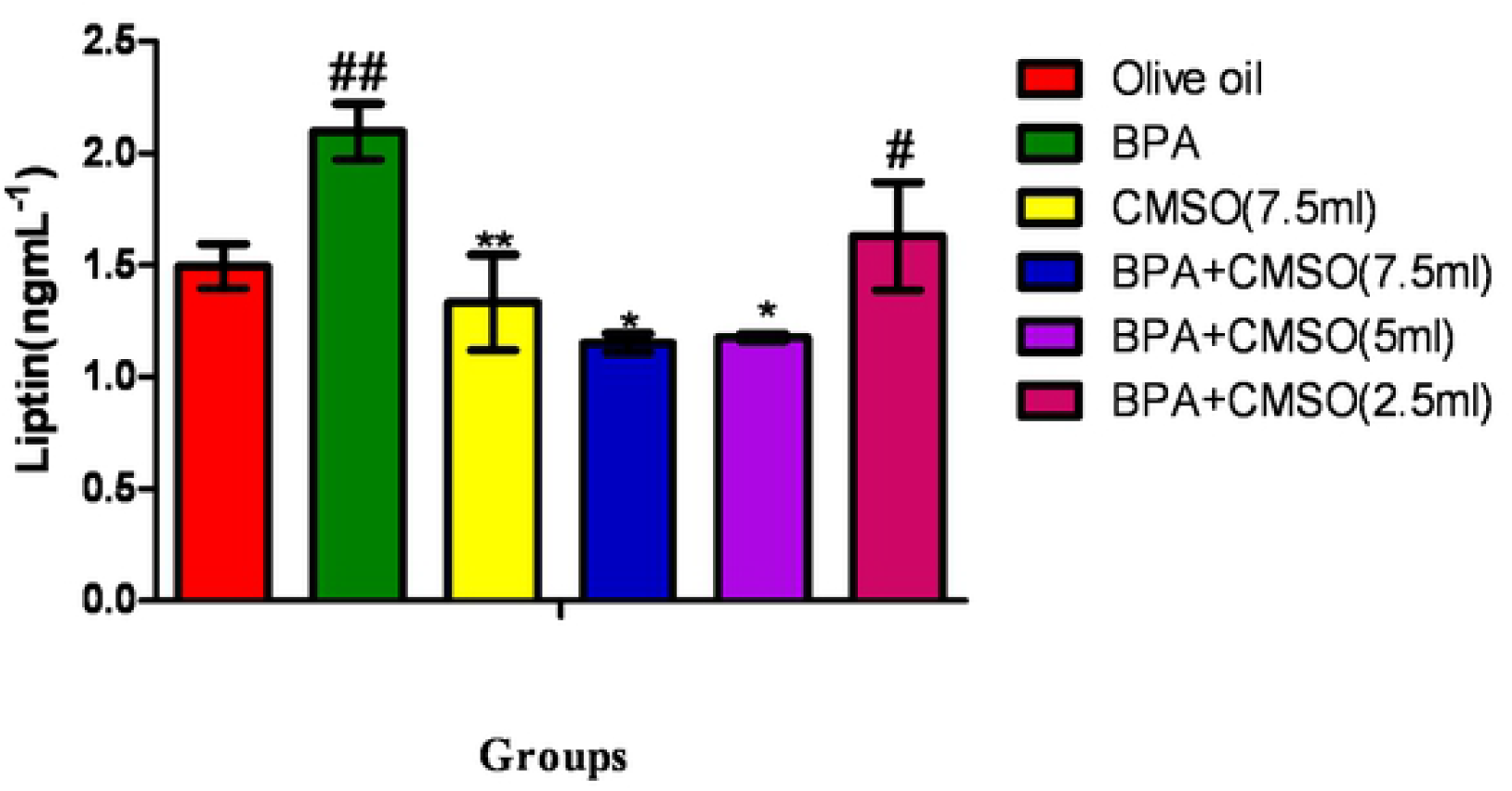
Effect of CMSO on Liptin level in adipose tissue in BPA interference in lipid metabolism in albino rats. Data are shown as mean ± S.D (n=6). Mean values with the different signs are significantly different at P<0.05. BPA (Bisphenol A), CMSO (*Cucumeropsis mannii* Seed Oil).

**Figure 14:**
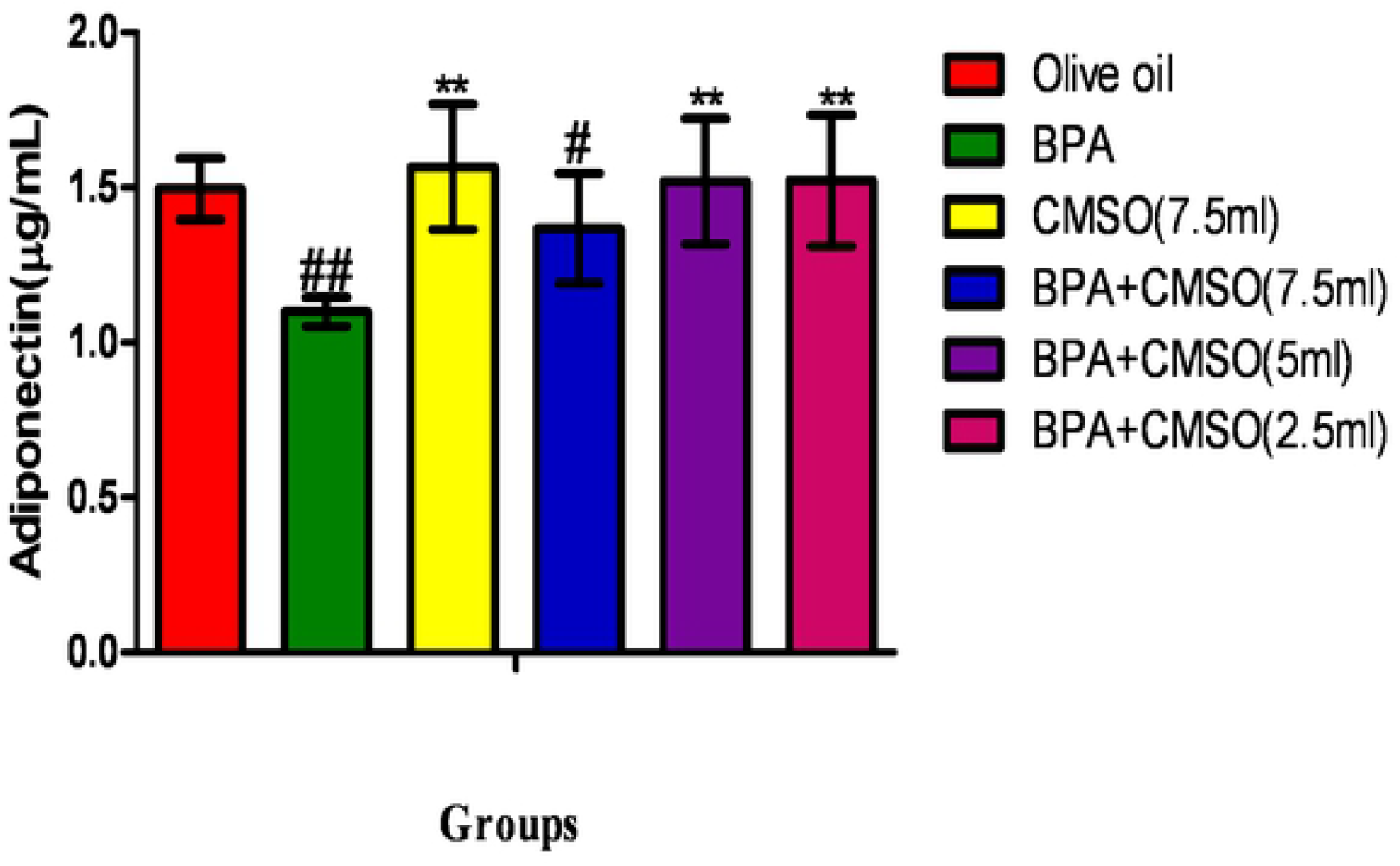
Effect of CMSO on Adiponectin level in adipose tissue in BPA interference in lipid metabolism in albino rats. Data are shown as mean ± S.D (n=6). Mean values with the different signs are significantly different at P<0.05. BPA (Bisphenol A), CMSO (*Cucumeropsis mannii* Seed Oil).

### 3.6 Effect of CMSO on Serum Liptin and Adiponectin levels in BPA interference of lipid metabolism in male rats

Bisphenol-A (BPA) administration significantly (p<0.05) elevated liptin level and reduced adiponectin level in male rats’ serum (Figures 15 and 16). Co-administration of BPA and CMSO in male rats significantly(p<0.05) reduced the level of liptin in a dose-dependent manner and elevated the adiponectin level in male rats as shown in Figures 15 and 16.

**Figure 15:**
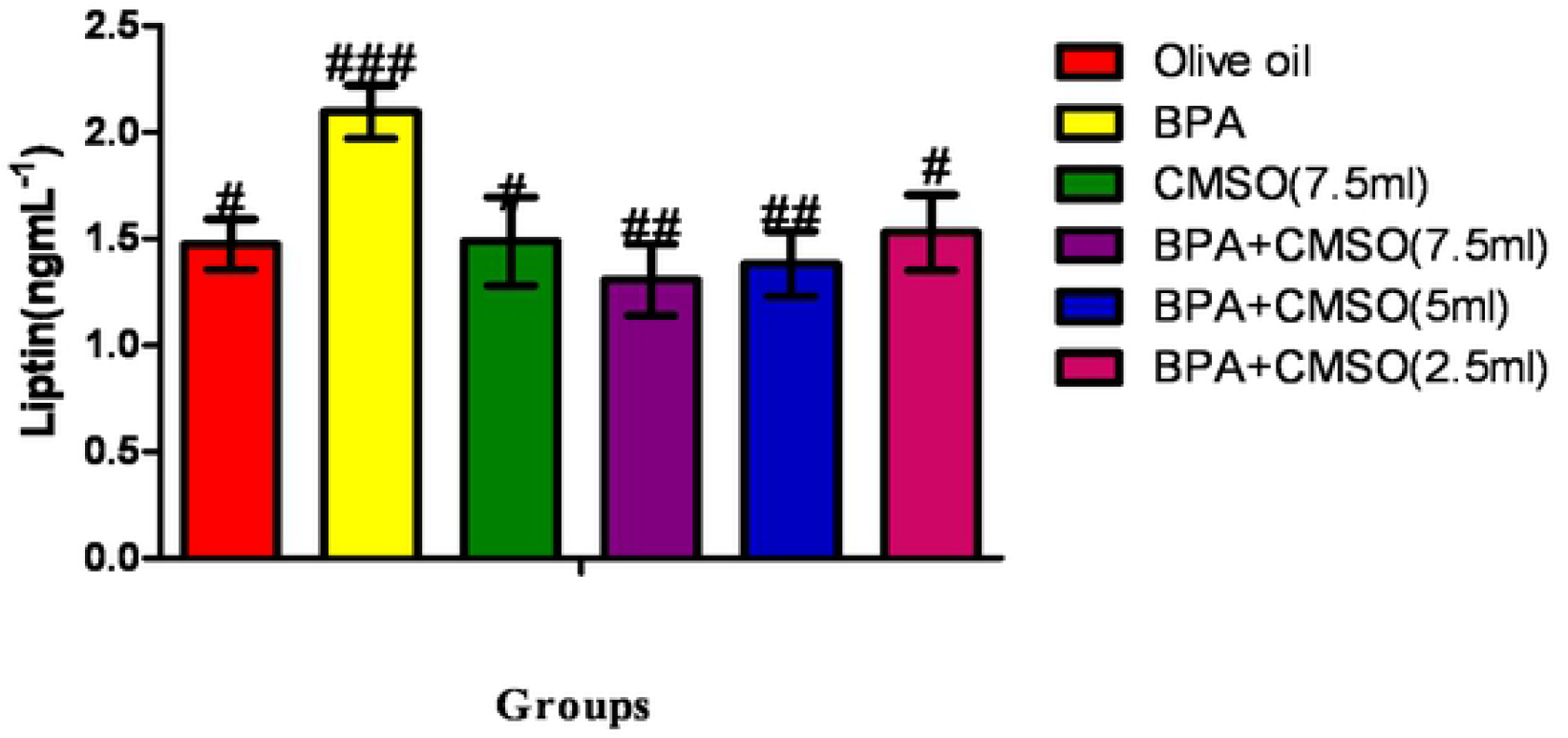
Effect of CMSO on Serum Liptin level in BPA interference in lipid metabolism in albino rats. Data are shown as mean ± S.D (n=6). Mean values with the different signs are significantly different at P<0.05. BPA (Bisphenol A), CMSO (*Cucumeropsis mannii* Seed Oil).

**Figure 16:**
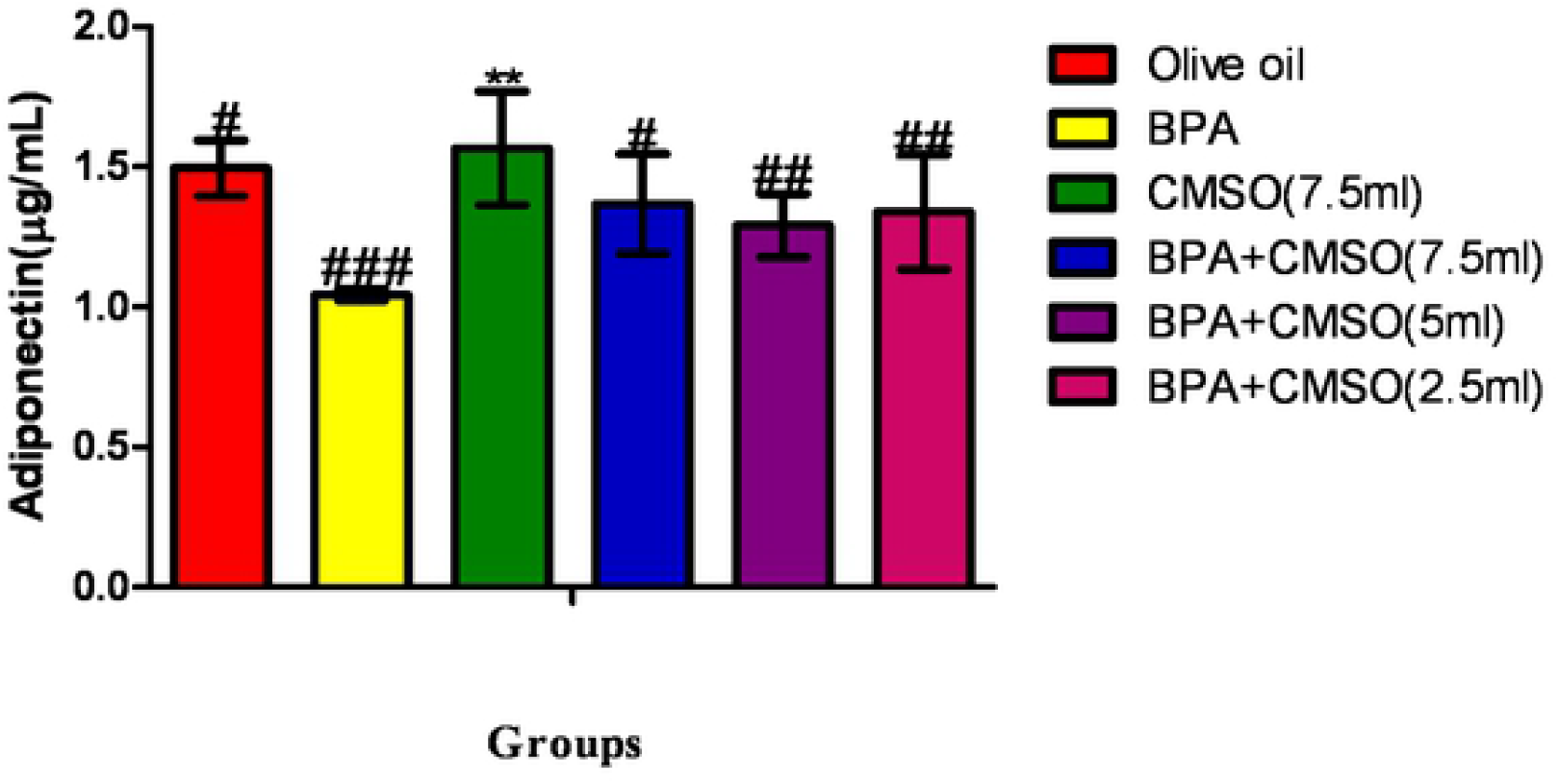
Effect of CMSO on Serum Adiponectin level in BPA interference in lipid metabolism in albino rats. Data are shown as mean ± S.D (n=6). Mean values with the different signs are significantly different at P<0.05. BPA (Bisphenol A), CMSO (*Cucumeropsis mannii* Seed Oil).

### 3.7 Effect of CMSO on Body weight of rats in BPA induced interference in lipid metabolism in rats

BPA administration in rats significantly (p<0.05) decreased the weights of the rats (Figure 17). However, there was a significant (p<0.05) increase in the bodyweight of rats after coadministration of BPA +CSMO (Figure 17).

**Figure 17:**
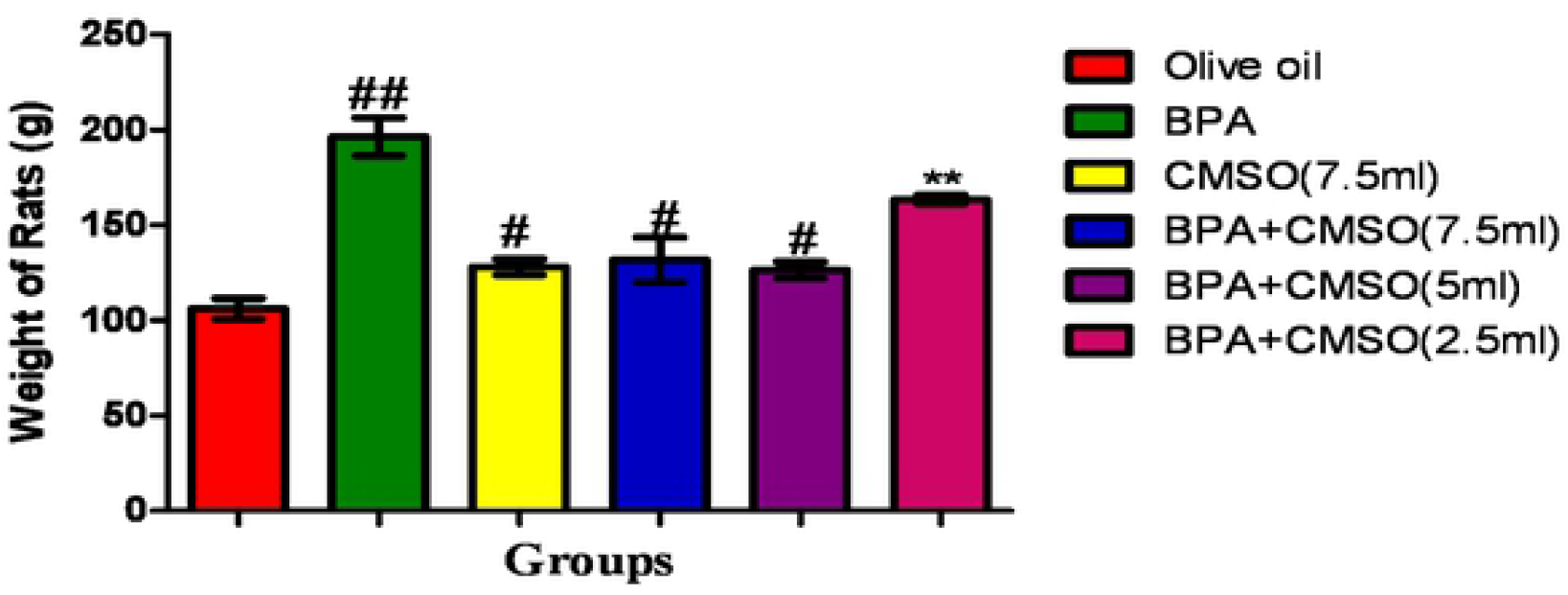
Effect of CMSO on Bodyweight in BPA interference in lipid metabolism in albino rats. Data are shown as mean ± S.D (n=6). BPA (Bisphenol-A), CMSO (*Cucumeropsis mannii* seed oil).

## 4. Discussion

### 4.1 Effect of CMSO on the Serum lipid profiles

In this present study, it was observed that administration of BPA significantly (p<0.05) elevated the serum and adipose tissues levels of cholesterol, triglycerides, and LDL-C with a reduction in HDL-C level in rats. These present findings are following the research carried out by National Center for Cardiovascular Diseases, (2016) that BPA exerted a drastic decrease in total cholesterol (TC), high-density lipoprotein cholesterol (HDL-C), with a deleterious increase in low-density lipoprotein cholesterol (LDL-C), triglyceride (TAG), lactate dehydrogenase (LDH) and creatine kinase-MB (CK-MB). Similarly, the present study is in line with a previous study (Zhu *et al*., 2015) that BPA caused a deleterious decrease in total cholesterol (TC), high-density lipoprotein cholesterol (HDL-C), with a rapid increase in low-density lipoprotein cholesterol (LDL-C), triglyceride (TG) and free fatty acids (FFA) thus altering the transport mechanism of HDL-C.

Interestingly, It has also been reported by Cai *et al*. (2015) that BPA conversely increased the level of and FFA in the blood with a corresponding increase in low-density lipoprotein (LDL-C) and triglycerides (TGs) in the adipose tissues of rats (Cai *et al*., 2015). The accumulation of LDL-C and TGs has been implicated in atherosclerosis (oxidized LDL-C within the walls of arteries) and obesity (Cai *et al*., 2015).

In furtherance to this, Gao *et al*. (2017) reported the hyperlipidemic and compounding effect of BPA in the induction of oxidative stress in serum and adipose tissues in rats that ultimately leads to the generation of ROS such as hydroxyl radical, peroxynitrite anion, nitric oxide, singlet oxygen, and the peroxyl radical thus, affirming its deleterious effect in this present study.

In this present study, the observed reduction in the level of HDL-C could be due to accumulation of metabolites and binding of the toxicants to the active site of the enzyme thus, decreasing the synthesis of3-hydroxy-3-methyl-glutaryl-coenzyme-A reductase (HMG-CoA reductase), an enzyme that catalyzes the conversion of 3-hydroxy-3-methyl-glutaryl-coenzyme-A to mevalonic acid, a necessary step in the biosynthesis of cholesterol or the reduction in the level of HDL-C could be due to BPA interference with apolipoprotein A1 (ApoA1), a protein that has a specific role in the transport and metabolism of HDL-C (the “good cholesterol) while increase in Cholesterol, LDL-C and TAGs could be due to BPA interference with apolipoprotein B (ApoB) (a protein that has a specific role in the transport of LDL-C) and the binding of the toxicants to the active site of lipase enzymes (an enzyme that breaks down triglycerides into free fatty acids and glycerol) that ultimately leads to hyperlipidemia and accumulation of cholesterol in the serum and adipose tissues.

However, co-administration of BPA and CMSO significantly (p<0.05) reduced cholesterol, triglycerides, and LDL-C with an elevation of HDL-C. The reduction in the level of cholesterol, triglycerides, and LDL-C with the elevation of HDL-C in serum and adipose tissues may be due to the antioxidant and chemical constituents of CMSO such as omega-6 fatty acids, omega-3 fatty acids, palmitic acids, stearic acids, oleic acids, linoleic acids and other monounsaturated and polyunsaturated fatty acids which has been reported to protect the functional and structural integrity of cell membrane as well as maintenance of membrane phospholipids that ultimately restored the activities of lipoprotein lipase and 3-hydroxy-3-methyl-glutaryl-coenzyme-A reductase (HMG-CoA reductase) (an enzyme involved in the breakdown of triglycerides into free fatty acids and glycerol and the biosynthesis of cholesterol respectively). Therefore, when hormones signal the need for energy, fatty acids and glycerol are released from triglycerides stored in fat cells (adipocytes) and are delivered to organs and tissues in the body or in times of stress when the body requires energy, fatty acids are released from adipose cells and mobilized for use (Iwaniec *et al*., 2007). The process begins when levels of glucagon and adrenaline in the blood increase and these hormones bind to specific receptors on the surface of adipose cells (Goliasch *et al*., 2015). This binding action starts a cascade of reactions in the cell that results in the activation of lipase that hydrolyzes triglyceride in the fat droplet to produce free fatty acids (Cai *et al*., 2015). These fatty acids are released into the circulatory system and delivered to skeletal and heart muscle as well as to the liver (Iwaniec *et al*., 2007). In the blood the fatty acids are bound to a protein called serum albumin; in muscle tissue, they are taken up by the cells and oxidized to carbon dioxide (CO_2_) and water to produce energy (Gao *et al*., 2017).

### 4.2 Effect of CMSO on the Liptin and adiponectin

In this present study, it was shown that BPA administration significantly (p<0.05) increased the levels of serum and adipose tissues of liptin with a concomitance decrease with serum and adipose tissues of adiponectin levels in rats. The results of this present study correlated with the investigation of Angle *et al*. (2013) who reported a drastic increase in serum liptin levels with a concomitance decrease in serum level of adiponectin that ultimately lead to abdominal fat mass. Similarly, It has been reported by Gao *et al*. (2008) that BPA induction increased the level of liptin in adipose tissues that ultimately affects the body weight acting mainly on the central nervous system and specifically the hypothalamus with a concomitance decrease in serum levels of adiponectin that plays a key role in the regulation of glucose uptake by the cells (Ben-Jonathan *et al*., 2009)

Interestingly, Valentino *et al*. (2013) carried out an *in vitro* study using human adipocytes and revealed that low doses of BPA impaired insulin-stimulated glucose utilization and insulin signaling pathway thus dysregulating adipocytes function. They also found in male offspring higher circulating liptin and insulin levels with a corresponding reduction of serum adiponectin thus indicating clear evidence of metabolic dysfunction in white adipose tissue caused by BPA exposure (Bouret *et al*., 2015). Similar to this present study, BPA has been reported to increase the level of liptin in adipose tissues thus, resulting in obesity and inflammatory-related diseases, including hypertension, metabolic syndrome, and cardiovascular disease (Cirillo *et al*., 2008). The increase in liptin level in the blood is the result of the bioaccumulation of liptin via the modulation of genes that regulate its activities.

Therefore, when hormones signal the need for energy, fatty acids and glycerol are released from triglycerides stored in fat cells (adipocytes) and are delivered to organs and tissues in the body or in times of stress when the body requires energy, fatty acids are released from adipose cells and mobilized for use (Iwaniec *et al*., 2007). The process begins when levels of glucagon and adrenaline in the blood increase and these hormones bind to specific receptors on the surface of adipose cells (Goliasch *et al*., 2015). This binding action starts a cascade of reactions in the cell that results in the activation of lipase that hydrolyzes triglyceride in the droplet to produce free fatty acids (Cai *et al*., 2015). These fatty acids are released into the circulatory system and delivered to skeletal and heart muscle as well as to the liver cell (Iwaniec *et al*., 2007). In the blood, the fatty acids are bound to serum albumin, taken up by the cells, and oxidized to carbon dioxide (CO_2_) and water to produce energy (Gao *et al*., 2017). The liver takes up a large fraction of the fatty acids (Iwaniec *et al*., 2007). There they are in part resynthesized into triglycerides and are transported in VLDL lipoproteins to muscle and other tissues (Iwaniec *et al*., 2007). A fraction is also converted to small ketone molecules that are exported via the circulation to peripheral tissues, where they are metabolized to yield energy (Sirtori, 2015).

### 4.3 Effect of CMSO on the Coronary risk index and atherogenic index

In this present study, Bisphenol-A (BPA) administration significantly (p<0.05) elevated the coronary risk index and atherogenic index (AI) in serum and adipose tissue of male rats. This present study is by the previous study (Bouret *et al*., 2015) who reported that BPA induction caused an increase in the atherogenic index (AI) and coronary risk index (CRI) in serum and adipose tissues of rats which is strong biomarkers for predicting the risk of Coronary artery disease (CAD).

Interestingly, this finding also correlated with a cross-sectional study conducted in Iran, that BPA administration in rats conversely elevated AI and CRI in serum and adipose tissues above normal range and was negatively associated with body mass index (Iwaniec *et al*., 2007).

Similarly, this present work is in line with another prospective study in Turkis that BPA induction in rats inversely increased AI in both serum and adipose tissues of male rats that ultimately lead to cardiovascular disease such as atherosclerosis and arteriosclerosis (Bouret *et al*., 2015) thus, thickening or hardening the loss of elasticity of the walls of arteries that ultimately restricts the blood flow to one’s organs and tissues and lead to severe health risks.

The results of this study are similar to the research carried out by (NCCD, 2016) who evaluated the effect of BPA in rats and integrated two lipids indies (TAGs and HDL-C) to generate AI which can be considered as a novel and better biomarker for obesity (Iwaniec *et al*., 2007). AI is calculated as log_10_ (TG/HDL-C), constructed as a biomarker of plasma atherosclerosis, and has been proved to be significantly correlated with other important atherosclerosis indexes such as LDL-C size and small-dense LDL-C (Knight *et al*., 2009).

In this present study, the increase in AI and CRI could be due to the accumulation of metabolites and binding of toxicants to the active site of the enzyme thus, decreasing the activity of lipase that ultimately lead to the buildup of fatty plaques, cholesterol, and some other substances in and on the artery wall (Zhu *et al*., 2015).

However, administration of BPA and CMSO significantly (p<0.05) reduced coronary risk index and atherogenic index. The reduction in the level of AI and CRI in serum and adipose tissues of rats may be due to the antioxidant and chemical constituents of CMSO.

### 4.4 Effect of CMSO on the Bodyweight of male rats

In this present study, BPA significantly (p<0.05) decreased the bodyweight of rats. These findings correlated with the observation of Gurmeet *et al*. (2014) who reported that the bodyweight of rats after BPA exposure slightly decreased when compared to the control group. Similarly, Miao *et al*. (2008) reported a significant (p<0.01) decrease in body weight and testicular volume of rats when compared with the normal control. Hence, some previous studies reported that after administration of BPA the body weight of male rats did not show a significant difference at a low dose of the extract (Korkmaz *et al*., 2010; Norazit *et al*., 2012; Nanjappa and Akingbemi, 2012). It was noted that BPA intoxication decreased the bodyweight of the experimental rats by damaging the important molecules, such as proteins in the testis. However, CMSO co-treatment significantly (p<0.05) increased the bodyweight of rats which may be due to the therapeutic efficacy and protective potential of CMSO.

## 4. Conclusion

Based on the findings from this present study, BPA exposure increased the serum and adipose tissues levels of cholesterol, triglycerides, LDL-C, liptin, coronary risk index, and atherogenic index with a reduction in body weight, HDL-C, and adiponectin in rats. Interestingly, the administration of CMSO reduced the BPA-induced toxicity through the restoration of serum lipid profiles, adiponectin, liptin, and bodyweight of rats. The results of this present study show the potentials and the therapeutic role of CMSO as a potent antioxidant capable of ameliorating BPA-induced oxidative stress and perturbation of membrane in male rats.

